# Tup1 is Critical for Transcriptional Repression in Quiescence in *S. cerevisiae*

**DOI:** 10.1101/2022.08.10.503497

**Authors:** Thomas B. Bailey, Phaedra A. Whitty, Eric U. Selker, Jeffrey. N. McKnight, Laura E. McKnight

## Abstract

Upon glucose starvation, *S. cerevisiae* shows a dramatic alteration in transcription, resulting in wide-scale repression of most genes and activation of some others. This coincides with an arrest of cellular proliferation. A subset of such cells enters quiescence, a reversible non-dividing state. Here, we demonstrate that the conserved transcriptional corepressor Tup1 is critical for transcriptional repression after glucose depletion. We show that Tup1-Ssn6 binds new targets upon glucose depletion, where it remains as the cells enter the G0 phase of the cell cycle. In addition, we show that Tup1 represses a variety of glucose metabolism and transport genes. We explored how Tup1 mediated repression is accomplished and demonstrated that Tup1 coordinates with the Rpd3L complex to deacetylate H3K23. We found that Tup1 coordinates with Isw2 to affect nucleosome positions at glucose transporter HXT family genes during G0. Finally, microscopy revealed that a quarter of cells with a Tup1 deletion contain multiple DAPI puncta. Taken together, these findings demonstrate the role of Tup1 in transcriptional reprogramming in response to environmental cues leading to the quiescent state.

## Introduction

Cellular quiescence is a non-proliferative cell state that appears conserved across all life (Marescal & Cheeseman, 2020; Rittershaus et al., 2013; van Velthoven & Rando, 2019). Quiescent cells are distinct from other non-dividing cells, such as senescent and terminally differentiated cells, in that quiescent cells can return to a proliferative state if the right conditions are met. Quiescence is necessary for the survival of unicellular organisms in environments that are unfavorable for proliferation; in fact, most microbes in nature are quiescent (O’Farrell, 2011; Valcourt et al., 2012). In multicellular eukaryotes, quiescence plays a critical role in various cell types including stem cells, fibroblasts, and lymphocytes (Tümpel & Rudolph, 2019; van Velthoven & Rando, 2019; Yao, 2014). The inability to properly regulate quiescence can lead to cancer; additionally, cancer stem cells can become quiescent, leading to drug resistance and the ability to initiate relapse at a later time (Vallette et al., 2019). All quiescent cells have conserved characteristics including transcriptional repression, increased cell density, translational repression, and decreased cell metabolism (de Virgilio, 2012; Gray et al., 2004; Miles et al., 2021; Sun & Gresham, 2021). Many of the components that regulate quiescence are found throughout life and likely have conserved functions; however, some of the fundamental ways cells regulate entry, exit, and maintenance of quiescence are not fully understood (Yanagida, 2009).

*Saccharomyces cerevisiae* is a well-established model organism for studying cellular quiescence because it can reliably produce a population of quiescent cells (Gray et al., 2004; Sun & Gresham, 2021). Quiescence is notoriously difficult to study in multicellular organisms due to the lack of quiescence-specific biomarkers and because it is difficult to determine which cells are quiescent until they begin dividing again. *S. cerevisiae* can produce a large population of quiescent cells that can be isolated by density centrifugation (Allen et al., 2006). Additionally, budding yeast is capable of surviving mutations that would be lethal in higher order organisms, allowing study of critically important factors in the process of establishing quiescence (Miles et al., 2021; Sun & Gresham, 2021). Because many of the fundamental components that regulate the cell cycle are conserved between budding yeast and higher eukaryotes, we suspect that many players involved in cell cycle arrest are highly conserved.

Tup1-Ssn6 is a transcriptional corepressor complex that globally regulates many cell processes in *S. cerevisiae* (Malavé & Dent, 2006). Tup1 serves mostly as a repressor, although it can also induce gene expression in some contexts depending on its phosphorylation state (Conlan et al., 1999; Proft and Struhl, 2002). Tup1 forms a complex with Ssn6 in which 3-4 Tup1 molecules associate with each Ssn6 molecule (Varanasi et al., 1996). The structure of Tup1 contains an N-terminal helix, which facilitates oligomerization and interacts with histone tails, and a beta-propeller domain composed of seven WD40 domains, which interact with other proteins such as histone deacetylases (HDACs) and DNA binding factors (Sprague et al., 2000). While the detailed mechanism of repression by Tup1 is unknown, some relevant information is available. Tup1 is recruited by sequence-specific transcription factors to DNA (Hanlon et al., 2011), where it recruits histone deacetylases to generate hypoacetylated regions of chromatin, typically associated with transcriptional repression (Davie et al., 2003; Fleming et al., 2014; Watson et al., 2000; Wu et al., 2001). In addition to HDAC recruitment, Tup1 interacts directly with nucleosomes by binding histone H3 and H4 tails, preferentially to hypoacetylated tails, to stabilize +1 nucleosomes that have been repositioned by Isw2 (Chen et al., 2013; Li & Reese, 2001; Rizzo et al., 2011; Zhang & Reese, 2004b). Tup1 also interacts directly with the Mediator complex through its subunit Hrs1, potentially inhibiting transcription into gene bodies (Papamichos-Chronakis et al., 2000). Lastly, Tup1 is thought to influence transcription by blocking the recruitment of activating factors to chromatin (Parnell & Stillman, 2011; Wong & Struhl, 2011).

Homologues of Tup1-Ssn6 found in higher eukaryotes include Groucho in drosophila, Grg in mice, and TLE1 in humans (Jennings & Ish- Horowicz, 2008). While the beta-propeller domain is well conserved in both sequence and structure, the sequence of the N-terminus is poorly conserved; however, it appears to be structurally conserved in simulations (Jumper et al., 2021; Varadi et al., 2022). Cross-species interactions have been established, where TLE family proteins can pull down yeast Ssn6 from cellular lysate, suggesting strong conservation of structure and function (Flores-Saaib & Courey, 2000; Grbavec et al., 1999). Mice and flies exhibit severe developmental phenotypes in Groucho/TLE null or hypomorphic mutations (Agarwal et al., 2015). Additionally, in humans TLE1 is currently being studied as both a cancer biomarker and a potential cancer therapeutic drug target (Ogawa et al., 2019; Yuan et al., 2017).

During quiescence in yeast, it has been observed that nucleosomes are repositioned and histone tails are deacetylated (McKnight, Boerma, et al., 2015; McKnight, Breeden, et al., 2015). Tup1 plays a role in histone deacetylation and chromatin remodeling in other contexts (Davie et al., 2003; Zhang & Reese, 2004b); however, it is unclear whether Tup1 contributes to these processes in quiescence. Tup1 has been identified as a gene that is essential for viability of cells in G0 and is implicated in glucose repression in yeast, but there is no known molecular mechanism to explain its role in these processes (Lin et al., 2017; Papamichos-Chronakis et al., 2004; Reimand et al., 2012; Williams et al., 1991; Williams & Trumbly, 1990). Here, we demonstrate that Tup1 is essential for repressing target genes in stationary phase by interacting with Rpd3 and Isw2 to generate regions of repressed chromatin. We also show that cells lacking Tup1 display morphological defects in stationary phase, suggestive of a possible role for Tup1 in mitosis.

## Results

### Tup1 Relocalizes After Glucose Starvation

Because Tup1 is a master regulator of many processes in *S. cerevisiae*, we sought to investigate the consequences of glucose starvation on Tup1 localization across the genome. We performed ChIP-Seq analysis on a Myc-tagged Tup1 strain during log phase, diauxic shift, and stationary phase (**Figure 1A**). Ideally, we would work with purified quiescent cells, but due to the unusual clumping of Tup1 deletion strains (**Supplementary Figure 1**), which is not alleviated by the addition of EDTA, it was impossible to isolate pure populations of quiescent cells. We therefore performed ChIP for Tup1 in stationary phase cells so that direct comparisons could be made to Δtup1 strains. However, we do anticipate that the findings in stationary phase will translate to quiescence because we observed that a fraction of stationary cells in Tup1 knockout strains are viable and can re-enter the cell cycle when plated on glucose rich media. Additionally, we know that Tup1 knockouts have reduced viability in G0, indicating that Tup1 plays a critical role in quiescence.

**Figure 1.**
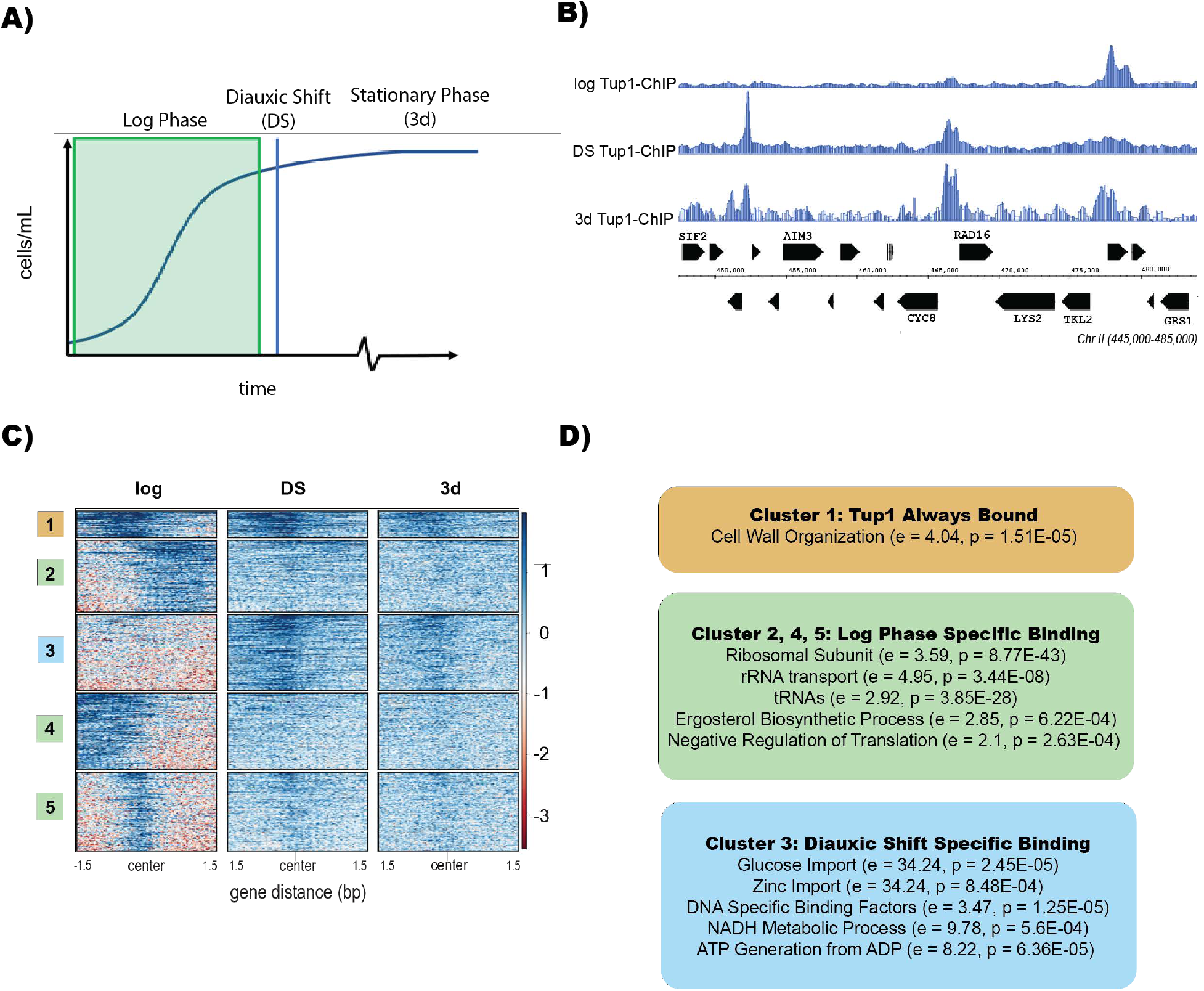
Tup1 Relocalizes After Glucose Starvation. **(A)** Illustration of a yeast growth curve indicating diauxic shift (DS) and stationary phase (3d). **(B)** Integrated Genome Browser (IGB) tracks representative of Tup1-myc ChIP-Seq during log phase, diauxic shift, and stationary phase. Data are normalized to RPKM. **(C)** ChIP-Seq heatmap of Tup1 binding during log phase, diauxic shift, and stationary phase. Heatmap was clustered using deeptools and sorted by binding intensity. **(D)** Gene ontology analysis from GOrilla of Tup1 binding locations with relative enrichment (e) and p-value (p) indicated for each term.

In log phase, we observed Tup1 binding to the promoters of roughly 923 genes, including genes for ribosomal protein subunits, tRNAs, and cell wall organization (**Figure 1B-C**). During diauxic shift, we observed Tup1 at 174 new targets across the genome, where it persisted into stationary phase. Gene ontology analysis revealed that during diauxic shift, Tup1 bound to the promoters of sugar transmembrane transporters, carbohydrate kinases, ATP from ADP factors, zinc transporters, and DNA binding transcription factors (**Figure 1D**). Motif analysis revealed significant enrichment for binding motifs of 78 transcription factors, including Mig1, which is known to recruit Tup1 to sites of repression (**Supplementary Table 1**) (Bailey & Grant, 2021; Papamichos-Chronakis et al., 2004; Treitel & Carlson, 1995).

### Tup1 Both Activates and Represses Key Targets for Quiescence Initiation

Next, we sought to understand how Tup1 binding correlates with transcriptional changes upon glucose repression. We performed differential expression analysis on RNA-Seq datasets, comparing transcription between wild type and Tup1 knockout strains during diauxic shift and stationary phase (**Figure 2A**). We were able to detect differential expression of 2102 genes in diauxic shift and 2999 genes during stationary phase in the Tup1 deletion, suggestive of a role for Tup1 in altering transcriptional programming during glucose exhaustion. Of the genes that were significantly differentially expressed in diauxic shift, nearly 60% had at least a 2-fold increase in expression, while the remaining 40% showed significant reduction in expression (**Figure 2A**). During stationary phase, only slightly more genes showed increased expression than showed decreased expression (approximately 52% versus 48%). Among genes that are repressed by Tup1, 787 were repressed in both diauxic shift and stationary phase (**Figure 2B**).

**Figure 2.**
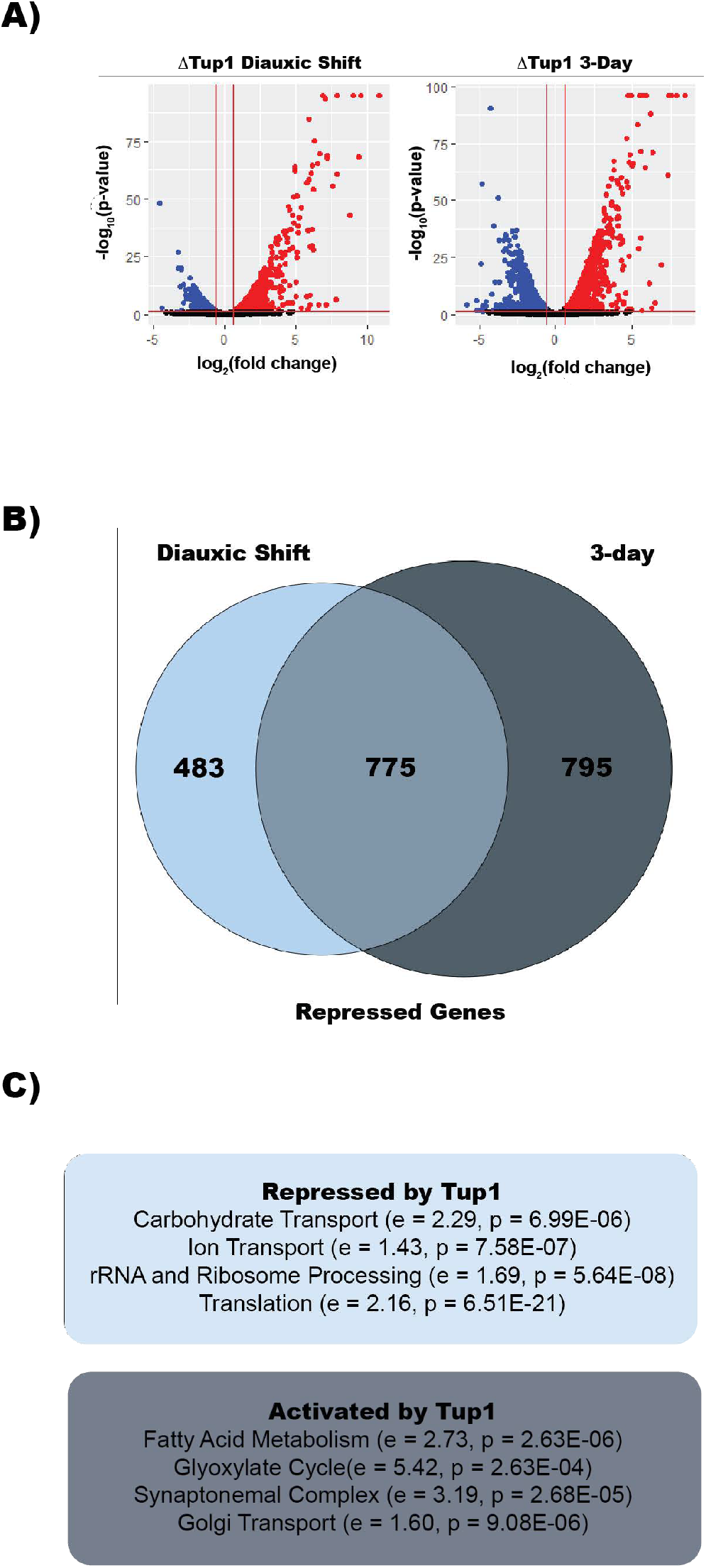
Tup1 Activates and Represses Key Targets for Quiescence Initiation. **(A)** Volcano plots of RNA-Seq differential expression data comparing a Tup1 deletion to wild type. Expression changes are based on two biological replicates. **(B)** Venn diagram comparing genes that are repressed by Tup1 in diauxic shift to those repressed in stationary phase. **(C)** Gene ontology analysis from GOrilla of transcripts affected by Tup1 in both diauxic shift and stationary phase with relative enrichment (e) and p-value (p) indicated for each term.

A previous screen for genes required for quiescence in yeast found 137 deletion mutants that were unable to form a quiescent cell population (L. Li et al., 2015). In our experiments, Tup1 was found to regulate expression of 57% of these genes, indicating that Tup1 plays a key role in transcriptional reprogramming during quiescence. Gene ontology analysis revealed that during diauxic shift, Tup1 represses genes associated with transmembrane transport and metabolism of carbohydrates (**Figure 2C**). We also observed Tup1 repression of genes implicated in transportation of amino acids, metal ions (Zn^2+^, Mg^2+^, Cu^2+^, and Fe^2+^), vitamins, and fatty acids. While transportation of fatty acids appeared repressed, metabolism of fatty acids appears to be activated by Tup1. In addition to repressing transportation of many components, Tup1 apparently stalls translation by repressing expression of ribosomal protein subunits, ribosome biogenesis genes, and tRNA ligases. During stationary phase, Tup1 represses many of the same genes found in diauxic shift, and also represses additional gene sets including rRNAs and DNA transcription factors associated with gene activation.

By comparing the results from RNA-Seq and ChIP-Seq, we found that Tup1 occupancy does not correlate solely with repression or activation of genes. For example, out of the 981 genes Tup1 releases from in DS, 35% have higher levels of expression in the Tup1 deletion and 25% have lower levels, while 40% are not significantly affected. Perhaps Tup1 release frees up binding for additional factors, which may be repressive or may induce transcription. Of the 176 genes where we detect Tup1 binding during DS, we find that 131 are differentially expressed. Most (60%) are repressed by Tup1 and less than 15% are activated, demonstrating that Tup1 recruitment to a site during diauxic shift is usually indicative of repression. Because the number of sites bound by Tup1 in the ChIP dataset does not match the number of repressed genes, it is likely that we did not detect all Tup1 binding that occurs during diauxic shift; however, we do not believe that the inability to detect all instances of Tup1 binding fully accounts for this difference. We observed that Tup1 binds to and represses many DNA binding transcription factors including Hap1, Nrg1, and Mig1 (**Supplementary Table 1**) and downstream effects of the repression of these transcription factors may account for why there are more repressed targets in the RNA-Seq dataset compared to the ChIP-Seq dataset.

### Sds3 and Xbp1 Repress Targets in Common With Tup1

Tup1 recruits the Rpd3L complex to sites of repression through DNA-bound transcription factors (Davie et al., 2003; Hanlon et al., 2011; Treitel & Carlson, 1995; Watson et al., 2000). We know that histone deacetylation during quiescence is largely Rpd3L-dependent and that Tup1 interacts with Rpd3L (McKnight, Boerma, et al., 2015). To explore this connection further, we first compared our Tup1 ChIP data to an existing Rpd3 ChIP dataset in quiescent cells (**Supplementary Figure 2A**) (McKnight, Boerma, et al., 2015). We found substantial overlap in Rpd3 and Tup1 peaks, indicating that they localize to similar targets, if not interact directly. Comparing sites of Tup1 binding in 3-day cultures to Rpd3 binding sites in purified quiescent cells (McKnight, Boerma, et al., 2015), we find that 160 of the 193 Tup1 peaks (94%) overlap with those of Rpd3. We also performed ChIP-Seq for the stress response factor Xbp1, which is required to recruit Rpd3 at Xbp1 motifs, and compared these data to Tup1 ChIP and found that Xbp1 and Tup1 also share binding sites (**Supplementary Figure 2B**).

To elucidate the relationship between Tup1 and Rpd3 in quiescence, we knocked out Sds3, an Rpd3L-specific subunit required for deacetylation, and compared RNA-Seq datasets to that of the Tup1 knockout (**Figure 3A**). Upon deletion of Sds3 we observed that 1680 genes were differentially expressed in diauxic shift (57% repressed by Sds3 and 43% activated) and 3212 are differentially expressed during stationary phase (52% were repressed by Sds3 and 48% were activated). 734 genes that were upregulated upon Sds3 deletion in diauxic shift were also found to be upregulated during the stationary phase. A gene ontology analysis revealed that Sds3 represses many genes in common with Tup1, including those implicated in carbohydrate transport and ribosome biogenesis. Sds3 is different from Tup1 in that it also represses cell cycle-associated genes.

**Figure 3.**
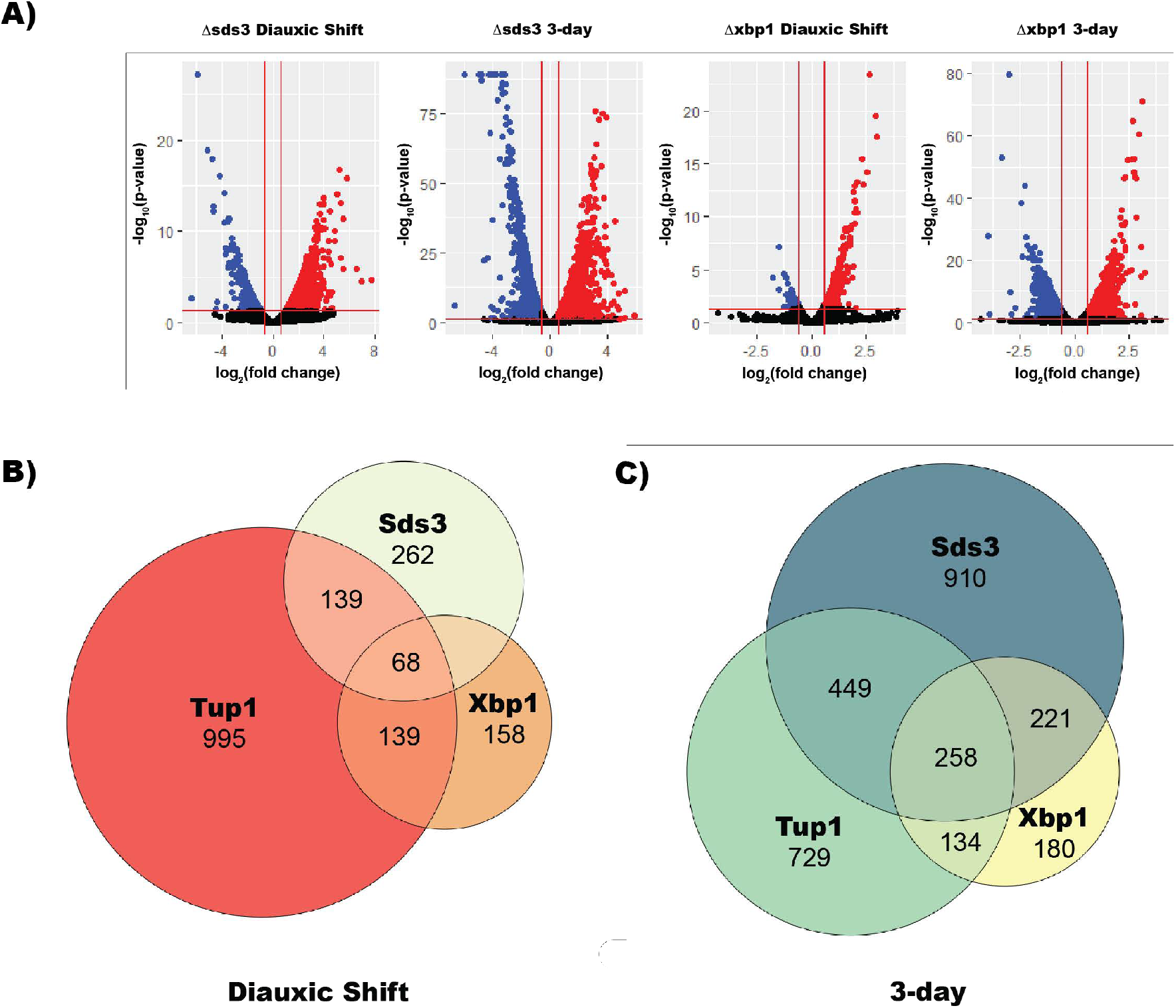
Sds3 and Xbp1 Repress Targets in Common with Tup1. **(A)** Volcano plots of RNA-Seq differential expression data comparing Xbp1 and Sds3 deletions to wild type in both diauxic shift and stationary phase. Expression changes are based on two biological replicates. **(B)** Venn diagram of transcripts repressed by Tup1, Sds3, and Xbp1 in diauxic shift. **(C)** Venn diagram of transcripts repressed by Tup1, Sds3, and Xbp1 in stationary phase.

When comparing the two RNA-Seq datasets we found that during diauxic shift, 37% of genes repressed by Sds3 were also repressed by Tup1 (p < 5.5e-42) (**Figure 3B**). During stationary phase, Tup1 represses 41% of genes repressed by Sds3 (p < 1.6e-65). This suggests widespread collaboration between Tup1 and Rpd3L in repressing target genes upon glucose starvation.

The transcriptional repressor Xbp1 has been shown to recruit Rpd3L to its binding sites (McKnight, Boerma, et al., 2015). MACS2 (Y. Zhang et al., 2008) peak calling from our ChIP dataset on Xbp1 and Tup1 binding suggests that Tup1 and Xbp1 localize to similar genes (Supplementary Figure 1). We performed ChIP-Seq experiments in strains lacking Xbp1 and found loss of Xbp1 has no effect on Tup1 binding, demonstrating that Xbp1 does not directly recruit Tup1, even though they colocalize (**Supplementary Figure 2B**). To measure the effect of Xbp1 deletion on transcriptional silencing, we deleted Xbp1 and performed RNA-Seq experiments during diauxic shift and log phase. We observed that in an Xbp1 deletion 528 genes are differentially expressed during diauxic shift, and during stationary phase 1437 genes were differentially expressed (**Figure 3A**).

Of these genes, 86% were repressed by Xbp1, and 14% were activated. During stationary phase, 51% of differentially expressed genes were repressed by Xbp1 and 49% activated. A gene ontology analysis revealed that Xbp1 is responsible for repressing genes associated with carbohydrate metabolism during diauxic shift, while repressing carbohydrate transport and cell cycle genes during stationary phase. Xbp1 and Tup1 repress 150 (p < 3.8e-28) shared genes during diauxic shift and 379 (p < 2.8e-60) genes during stationary phase (**Figure 3C**). Xbp1 represses 153 (p < 4.8e-45) genes in common with Sds3 during diauxic shift and 455 (p <1.3e-101) during stationary phase. Xbp1 also regulates cell cycle genes along with Sds3, while Tup1 has no effect on these genes.

### Tup1, Sds3, and Xbp1 Are Required for H3K23 Deacetylation at Repressed Genes

Since Tup1 facilitates H3K23 deacetylation in other contexts (Malavé & Dent, 2006), we next wanted to see if Tup1 contributes to the H3K23 deacetylation we observe during quiescence. To determine if Tup1 is required for deacetylation, we performed H3K23ac ChIP-Seq during stationary phase where Tup1, Xbp1, or Sds3 had been deleted. More than half of the genes (535) with changes in acetylation showed differential regulation by Tup1 or Sds3 in our RNA-seq data (p < 7.97e-6). Cells with the Sds3 deletion showed global hyperacetylation in the promoter region during the stationary phase (**Figure 4A-B**). Additionally, Δtup1 cells also showed global hyperacetylation, though not as dramatically as with the Δsds3 strain. It was difficult to see a global effect of Xbp1 deletion on acetylation levels; however, when we filtered genes with an Xbp1 motif, it appears that deletion of Xbp1 influences acetylation levels as well, although not nearly as dramatically as with loss of Sds3 or Tup1. To determine the effect of acetylation of transcription, we took the 1000 genes with the greatest changes in acetylation levels across all three mutants and compared them to genes repressed by Tup1 or Sds3 in stationary phase (**Figure 4C**). We observed a strong relationship between genes that are deacetylated in stationary phase and genes that are repressed.

**Figure 4.**
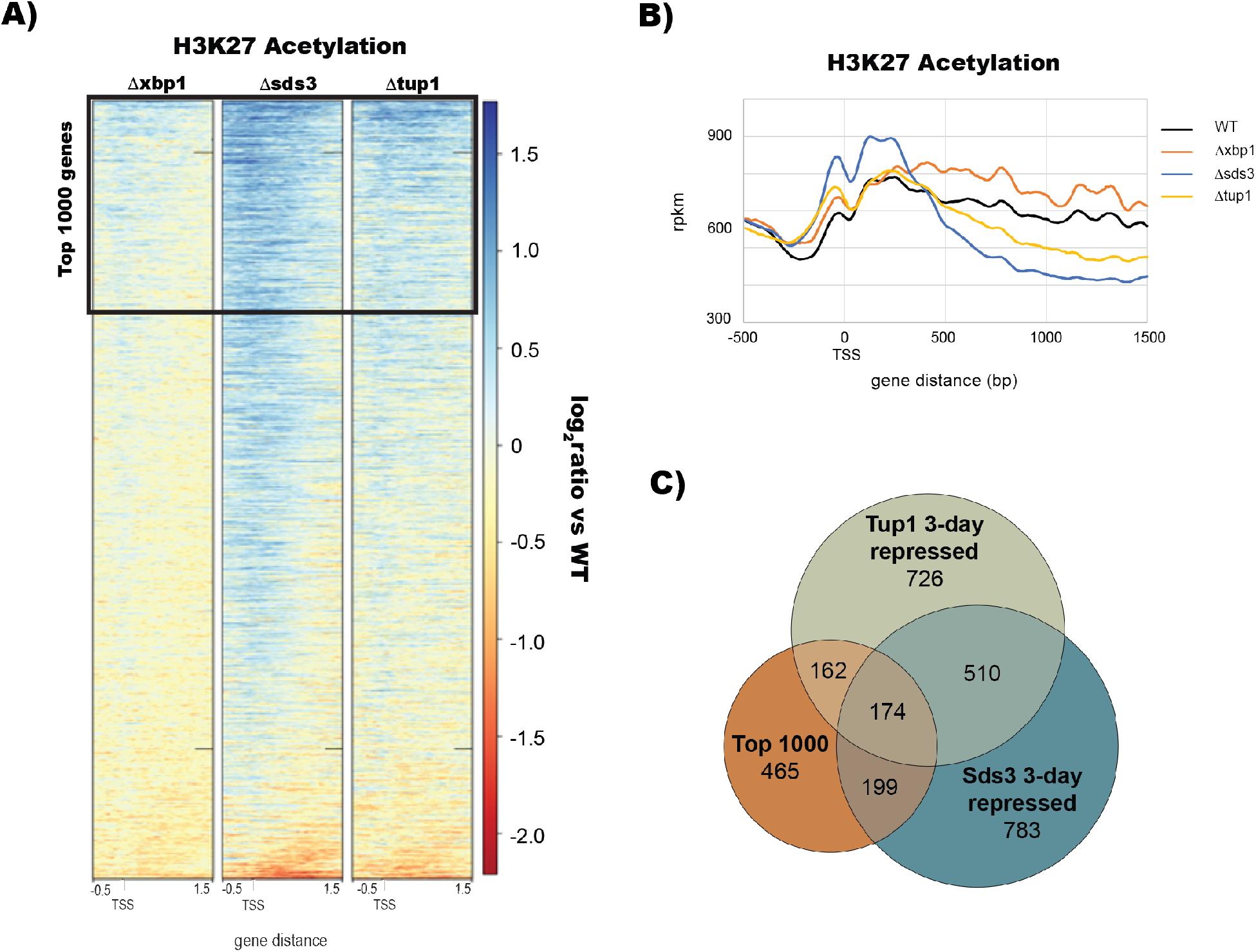
Tup1, Sds3, and Xbp1 are Required for H3K23 Deacetylation at Repressed Genes. **(A)** Heatmap showing the change in H3K23ac levels in deletion mutants compared to wild-type across all yeast genes. **(B)** H3K23ac profile levels across gene promoters, integrated across all yeast genes. **(C)** Venn diagram comparing the 1000 genes with the largest change in H3K23ac to genes repressed by Tup1 and Sds3 in stationary phase.

### Tup1 and Isw2 Regulate Nucleosome Positions at HXT Family Genes

During quiescence, cells reposition their +1 nucleosomes, either to shrink the nucleosome depleted region (NDR) or to expand it. Shrinking of the NDR is generally regarded as repressive, because it presumably blocks recruitment of DNA transcription factors and RNA polymerase II. Expansion of the nucleosome NDR, on the other hand, presumably facilitates recruitment of RNA polymerase and is associated with active transcription. The vast majority of NDRs shrink during quiescence in yeast, while a few are expanded (McKnight, Breeden, et al., 2015). The chromatin remodeler Isw2 is known to interact with Tup1, and both proteins are required to position nucleosomes in some contexts (Rizzo et al., 2011; Zhang & Reese, 2004b, 2004a). Because Tup1 is involved in +1 nucleosome stabilization in other contexts, we performed MNase- Seq to determine if Tup1 affects the position of nucleosomes in stationary phase. We found that specifically, Tup1 and Isw2 are required to position nucleosomes at the HXT family of glucose transporter genes HXT1, HXT3, HXT8, HXT9, HXT11, HXT13, HXT15, and HXT16 (**Figure 5A**).

**Figure 5.**
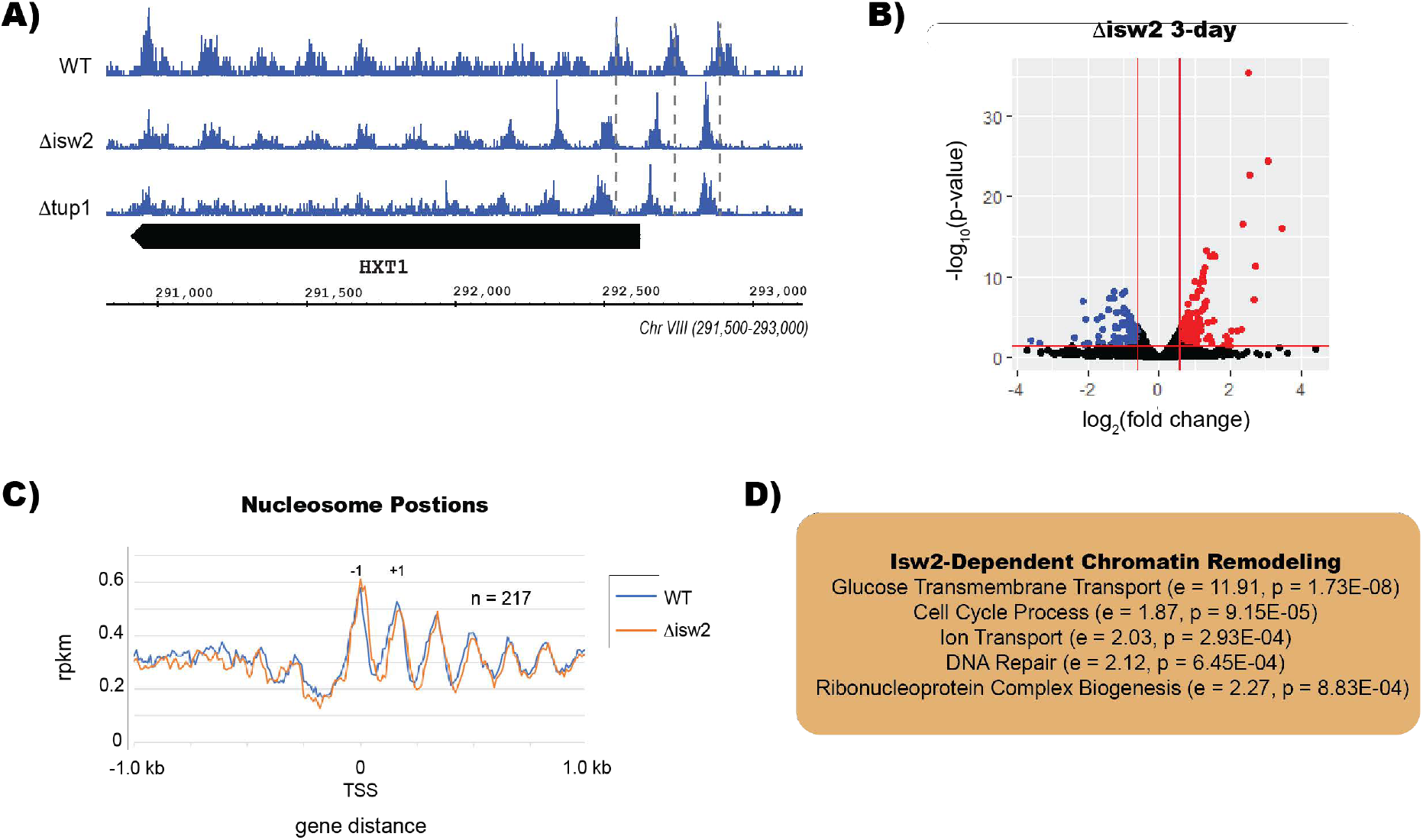
Tup1 and Isw2 Regulate Nucleosome Positions at HXT Family Genes. **(A)** Integrated Genome Browser (IGB) tracks of MNase-Seq in wild type, Δisw2, and Δtup1 in stationary phase. Dashed gray lines indicate wild type nucleosome positions. **(B)** Volcano plot of Δisw2 deletion in stationary phase demonstrating shifted nucleosomes at the HXT1 locus. **(C)** MNase-Seq dyad signal across 217 genes with shifted nucleosomes in Δisw2 during stationary phase. “-1” indicates the first nucleosome upstream of the TSS, while “+1” indicates the first nucleosome downstream of the TSS. **(D)** Gene ontology terms for genes with shifted nucleosome positions in Δisw2 yeast at stationary phase, with relative enrichment (e) and p- value (p) indicated for each term.

RNA-Seq data revealed that Isw2 is required to repress 106 genes during diauxic shift and 172 genes in stationary phase (**Figure 5B**). Isw2 represses many shared targets with Tup1: 33% of genes repressed by Isw2 are also repressed by Tup1 in diauxic shift, whereas almost half of the genes repressed by Isw2 in stationary phase are repressed by Tup1. In addition, we found a greater number of mispositioned nucleosomes in the Δisw2 strain, which had mispositioned nucleosomes in the −2, −1, +1, and +2 positions relative to the transcription start site (TSS) (**Figure 5C**). Our gene ontology analysis of the genes with mis-positioned nucleosomes in the Δisw2 deletion showed that Isw2 regulates nucleosomes in the promoters of genes involved in glucose transport, cell cycle, and DNA repair pathways in stationary phase (**Figure 5D**).

### Deletion of Tup1 or Sds3 Leads to Morphological Defects in Stationary Phase

Due to the multimeric nature of the Tup1-Ssn6 complex, it has been hypothesized that Tup1 may act as a scaffold for long-distance chromatin interactions. We know that during quiescence there are large scale chromatin folding events in yeast (Swygert et al., 2019). We have biochemical evidence that Tup1 may regulate chromatin folding in vitro; however, it is unknown whether this occurs in vivo. We initially wondered if deleting Tup1 would lead to less compaction of chromatin and sought to determine if Δtup1 cells had larger nuclei using fluorescence microscopy and DAPI staining. Although we failed to detect statistically significant changes in nuclei size, we found that Δtup1 cells showed multiple DAPI puncta not observed in wild type cells (**Figure 6A-B**).

**Figure 6.**
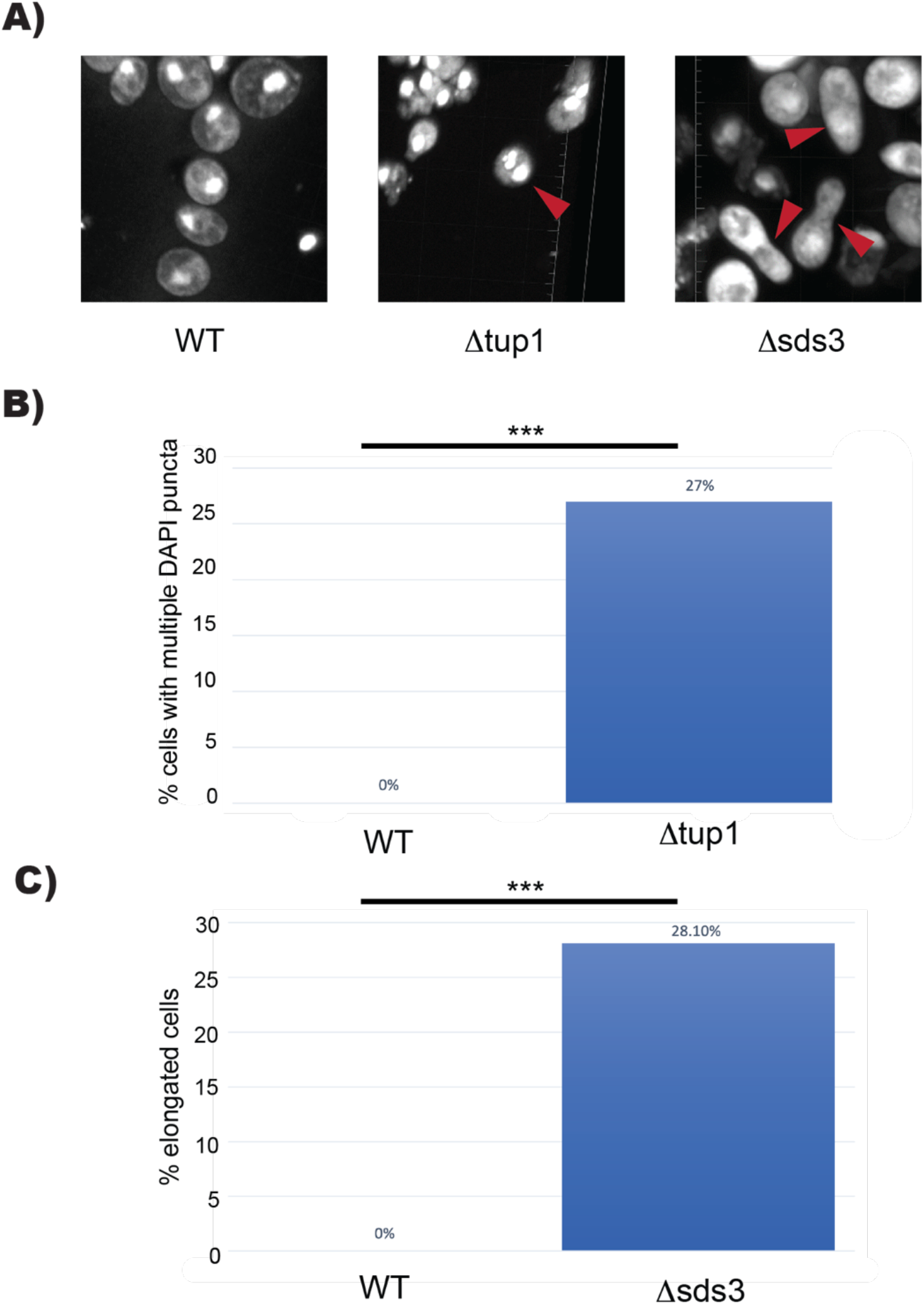
Deletion of Tup1 or Sds3 Leads to Morphology Defects in Stationary Phase. **(A)** Fluorescence microscopy of DAPI stained nuclei across the genome at 67x magnification in wild type, Δtup1, and Δsds3 yeast. **(B)** Quantification of Δtup1 cells with multiple DAPI puncta. WT n=100, Δtup1 n=48. **(C)** Quantification of peanut-shaped cells in Δsds3 yeast. WT n=100, Δsds3 n=64. *** p < 0.05

Deletion of Sds3 did not cause this phenotype; however, Δsds3 cells were often misshapen, which we termed “peanut shaped cells” (**Figure 6C**). Peanut shaped cells have been observed in other mutants, particularly mutants involved in schmoo formation such as AFR1, PEA2, and SPA2 (Chenevert et al., 1994). While Sds3 does not appear to regulate the expression of these genes to a significant extent, Sds3 is involved in regulating the cell cycle, bud site selection, and cell wall organization. While the phenotype between Δsds3 cells and Afr1 mutant appear similar, it is likely caused by different types of misregulation. We did not observe any changes in Δisw2 yeast compared to wild type yeast. The morphological defects observed in both Δtup1 and Δsds3 yeast suggest a potential mitotic defect, leading to aberrant distribution of DNA within nuclei in Δtup1 cells and abnormal cell division in the Δsds3 mutant. While we were unable to uncover the mechanism underlying this phenomenon, these findings may be of interest for further avenues of investigation.

## Discussion

Tup1 plays a key role in regulating large scale transcriptional responses to environmental conditions in yeast (Hanlon et al., 2011). Our ChIP-Seq studies with epitope-tagged Tup1 revealed that when glucose is depleted during the diauxic shift, Tup1 releases from numerous chromatin sites where it is bound in log phase and binds new positions that persist into the stationary phase. Tup1 binds some other sites in both log phase and during diauxic shift, typically associated with genes involved in cell wall organization. During log phase, Tup1 is bound to the promoters of ribosomal subunit proteins and tRNAs, which are actively transcribed at this stage but become repressed during diauxic shift; it seems that Tup1 is involved in both gene activation and repression. During diauxic shift, Tup1 relocalizes to genes associated with carbohydrate metabolism and transport, as well as factors that convert ADP to ATP, zinc transporters, and DNA-binding transcription factors; Tup1 relocalization to these genes represses their expression. The sites occupied by Tup1 regardless of glucose availability all become derepressed in a Tup1 knockout strain, indicating that Tup1 represses these targets. We suspect that repression of transcription factors by Tup1 may play a role in its ability to affect transcription of more factors in addition to those directly associated with its binding sites. Additionally, because Tup1 does not directly interact with DNA, we suspect that ChIP fails to reveal every instance of Tup1 binding.

Tup1 null mutants fail to repress many of the genes required for diauxic shift and establishment of quiescence. Many of the targets that are repressed/activated by Tup1 account for characteristics of quiescent cells. For example, common hallmarks of quiescent cells are fewer ribosomes, stalled translation, cell wall reorganization, and repression of metabolism. Our RNA-Seq data indicate that Tup1 is required to repress genes associated with all of these processes, demonstrating its role in regulating the transition to quiescence. Li et. al performed a genetic screen for factors that are required for quiescence in yeast. We found that Tup1 regulates more than 56% of these factors, further demonstrating the importance of Tup1 in regulating quiescence.

Tup1 regulates metabolism by simultaneously repressing genes involved in RNA, lipid, amino acid, and carbohydrate biosynthetic processes and activating the glyoxylate cycle and fatty acid metabolism. Transmembrane transporters of amino acids, carbohydrates, fatty acids, and metals (specifically magnesium, zinc, copper, and iron) are repressed by Tup1. We also find that two of the three yeast aquaporins are repressed by Tup1. It has been observed in other organisms that transmembrane transport is related to cellular proliferation (Peters-Regehr et al., 1999; Shyh-Chang & Daley, 2015). These findings indicate that yeast cells repress the vast majority of transmembrane transporters, which likely allows the cells to maintain a controlled intracellular environment in a context where metabolism is dramatically slowed.

The importance of the Rpd3 complex in establishing a quiescent state has been well demonstrated, and one factor that recruits Rpd3 is the transcription factor Xbp1 (McKnight, Boerma, et al., 2015). Xbp1 and Rpd3 share many common targets of repression with Tup1, while simultaneously having targets independent of Tup1. For example, Tup1 and Sds3 both strongly repress tRNA processing factors, ribosomal subunits, transmembrane transport, sugar catabolism, and cell wall components. Xbp1 and Tup1 overlap in their repression of transmembrane transport genes and sugar metabolism. However, Xbp1 and Sds3 strongly repress cell cycle genes, which are unaffected by Tup1.

Histone deacetylation is strongly associated with transcriptional repression in quiescence (McKnight, Breeden, et al., 2015) and we demonstrated the importance of Tup1, Xbp1, and Sds3 to facilitate histone deacetylation. Whether or not histone deacetylation is the main driving force of transcriptional repression is up for debate. We are only able to detect changes in histone deacetylation three days into stationary phase, which raises the possibility that histone deacetylation is not the main driving force in repressing transcription. However, recent findings demonstrate that the acetylome can change within 5 minutes of glucose depletion, so perhaps changes in the acetylome during diauxic shift are too subtle for us to detect (Hsieh et al., 2022; Cai, et al., 2011). We know that Tup1 and Rpd3 have repressive activities outside of histone deacetylation Malavé, & Dent, 2006), but it is difficult to decipher which mechanism is employed to repress targets. Overall, there is a strong correlation between genes that are repressed and those that lose their acetyl histone marks.

Tup1 and Isw2 both coordinately regulate nucleosome positions in the promoter regions of HXT family genes. Our RNA-Seq dataset revealed that these genes are also repressed by Tup1 and Isw2, suggesting that repositioning of nucleosomes by Isw2 and Tup1 at these genes leads to repression. Interestingly, these are the only genes in which both Tup1 and Isw2 affect nucleosome positions; deletion of Tup1 has little effect on the positions of nucleosomes at any other genes. We see a greater number of nucleosomes that are affected by Isw2 in genes involved in DNA repair pathways, glucose transport, and cell cycle process. We found that in a Sds3 knockout, cells form peanut shapes, indicative of a cell attempting to divide but ultimately failing to complete the process. Sds3 regulates many cell cycle genes, and it seems likely that failure to repress key cell cycle genes leads to the incomplete cell divisions we observed.

The multiple DAPI puncta observed in Tup1 knockout cells warrant further investigation. One possibility is that these cells have multiple nuclei or fragments of nuclei, which could be the result of a failure to properly segregate chromosomes during mitosis. Another possibility is these puncta are fragments of a broken-down nucleus, which is the result of cell cycle dysregulation. In any case, the observation highlights the importance of Tup1 in regulating the exit from the cell cycle. Taken together these results indicate that Tup1 is critical for gene repression in the exit from the cell cycle and establishment of quiescence.

It has been demonstrated that the Tup1 homologue GRG5 is critical for embryonic stem cell fate decisions in mice (Chanoumidou et al., 2018). Additionally, TLE1 is associated with many cancers in humans, including synovial carcinoma, lung cancer, breast cancer, glioblastoma, gastric cancer, and pancreatic cancer (Yuan et al., 2017). This indicates a role of Tup1/Groucho/TLE family proteins in determining cell fate in higher eukaryotes. Perhaps a conserved role of Tup1 family proteins is to regulate entry and exit from the cell cycle, which may explain why dysregulation of Groucho of TLE in higher eukaryotes leads to cancer.

## Materials and Methods

### Generation of Deletion Strains

All yeast strains were generated from the parent strain *S. cerevisiae* W303 RAD5+ (ATCC 208352). Null mutants were generated by replacing the gene of interest with antibiotic resistance markers amplified from pAG vectors (Longtine et al., 1998). FLAG- and Myc- tagged strains were made from a pFA6a vector containing their respective sequences. We then inserted the tags through homologous recombination of PCR products using selectable drug markers (Longtine et al., 1998).

### Growth Conditions

*S. cerevisiae* W303 cultures were streaked from glycerol stocks onto yeast extract peptone dextrose (YPD) plates and grown at 30° C. Individual colonies were picked and used to inoculate 25 mL YPD with shaking at 30° C. These cultures were grown to an O.D. of 0.6 to 0.8 for log phase cultures. Diauxic shift cultures were obtained by monitoring glucose levels with glucose test strips (Precision Laboratories). Cells were harvested two hours after there was no detectable glucose. Stationary phase cultures were prepared by growing yeast for three days in YPD medium.

### Chromatin Immunoprecipitation

Chromatin immunoprecipitation was performed as previously described (Rodriguez et al.; 201*4*). 100 OD-mL of cells were crosslinked in 1% formaldehyde for 20 minutes at 30° C. Cells were then quenched with a final concentration of 125 mM glycine, pelleted, and frozen at -80° C. Cells were lysed in a Bead Beater (Biospec) for two minutes with acid- washed glass beads, diameter *4*25-600 m (Sigma Aldrich) in the presence of protease inhibitors (Proteoloc Protease Inhibitor Cocktail, Expedeon) and then sheared in a Bioruptor sonicator for a total of 30 min (high output 30 sec on 30 sec off for 10 minutes 3×) and then centrifuged. The sheared chromatin supernatant was incubated with antibodies bound to G protein beads (Invitrogen). The crosslinks were reversed, overnight at 65° C and the sample was treated with 20 g RNase A for two hours at *4*5° C and 100 g Proteinase K for *4* hours at 55° C, and then purified using Qiagen MinElute columns. DNA samples were quantified using a Qubit fluorescence assay and libraries were prepared using a NuGEN Ovation Ultralow kit. Sequencing was performed by the University of Oregon Genomics and Cell Characterization Core Facility on an Illumina NextSeq500 on the 37 cycle, paired-end setting, yielding approximately 10-20 million paired- end reads per sample.

### RNA Extraction

Cells were removed from −80° C and ground with a mortar and pestle in liquid nitrogen. RNA was extracted using hot acid-phenol (Uppuluri et al., 2007) and cleaned up using the RNeasy kit (Qiagen). Libraries were prepared using the NuGEN Universal Plus mRNA kit. Sequencing was performed by University of Oregon Genomics and Cell Characterization Core Facility on an Illumina NextSeq500 with 37 cycles of paired-end setting. Paired end reads were filtered and aligned to the *S. cerevisiae* sacSer3 genome using bowtie2 (Langmead & Salzberg, 2012). Differential expression analysis was performed using DESeq2 (Love et al., 2014).

### Micrococcal Nuclease Digestion

We utilized our rapid MNase protocol (McKnight et al., 2021). In short, cross-linked cells were permeabilized by zymolyase digestion then treated with MNase and Exonuclease III. Next, the samples were treated with RNaseA and Proteinase K and purified using the Qiagen MinElute PCR Cleanup kit.

### DAPI Staining and Fluorescence Microscopy

Staining and microscopy was performed as described by Swygert et al. (2019). Cells were fixed in 3.7% formaldehyde for 20 min at 4° C in 0.1 M KPO4 pH 6.4 buffer. Cells were washed twice in sorbitol citrate (1.2 M sorbitol, 100 mM K2HPO4, 36.4 mM citric acid) and incubated with 2.5 μg of Zymolyase for 30 minutes. Cells were added to polytetrafluoroethylene (PFTE) slides with 0.1% polylysine then washed with ice cold methanol and acetone. Next 5 μL of DAPI mount (0.1 μg/ml DAPI, 9.25 mM p-phenylenediamine, dissolved in PBS, and 90% glycerol) was added to the well. Images were taken on a Zeiss LSM 880 confocal microscope with Airyscan. Images were analyzed and processed using Imaris software (Oxford Instruments). Two biological replicates were used for each strain.

### Computational Analysis

All alignments were performed using Bowtie2 (Langmead & Salzberg, 2012). Further analysis was performed using deepTools including generation of heatmaps (Ramírez et al., 2016). For ChIP, data was normalized to RPKM and gene tracks were visualized using Integrated Genome Browser (Freese et al., 2016). Peak picking from ChIP data was performed using MACS2 (Y. Zhang et al., 2008). Venn diagrams were generated using euler.com (Larson, 2020). Motif analysis was performed using SEA in MEME-suite (Bailey & Grant, 2021). Gene ontology was performed using Gorilla (Eden, 2015). We chose the two unranked lists option with our target set being genes that were present in our peak calling or genes that were differentially expressed in our RNA-seq datasets. Our background set included all transcripts that were detected in RNA-seq. Additionally we chose all enriched GO terms.

## Author Contributions

Conceptualization, T.BB., E.U.S., J.N.M.; Methodology, T.B.B, L.E.M., J.N.M.; Investigation, T.B.B., P.A.A., L.E.M., J.N.M.; Additional Resources, E.U.S.; Writing - Original Draft, T.B.B.; Writing - Reviewing and Editing, T.B.B., E.U.S.; L.E.M.; Visualization, T.B.B., L.E.M.; Supervision, E.U.S., J.N.M., L.E.M.; Project Administration, L.E.M., J.N.M.; Funding Acquisition, J.N.M.

## Declaration of Interest

The authors have no competing interests to declare.

## Data Availability

The datasets generated during this study are available at GEO under accession code GSE210934.

## Acknowledgments

The authors thank the Genomics Core (GC3F) at the University of Oregon for high throughput sequencing services. The authors also thank Christine Cucinotta (Fred Hutchinson Cancer Research Center) and Hideki Tanizawa, Osamu Iwasaki, Ken-Ichi Noma for helpful discussions related to this work. Additionally we thank Christine Cucinotta and Alison Greenlaw for tracking down Rpd3 data for us. This work was supported by a National Institutes of Health training grant T32 GM007759 (to T.B.B), and by NIGMS R01 GM129242 (to J.N.M. and L.E.M).

---

**Supplementary Figure 1.**
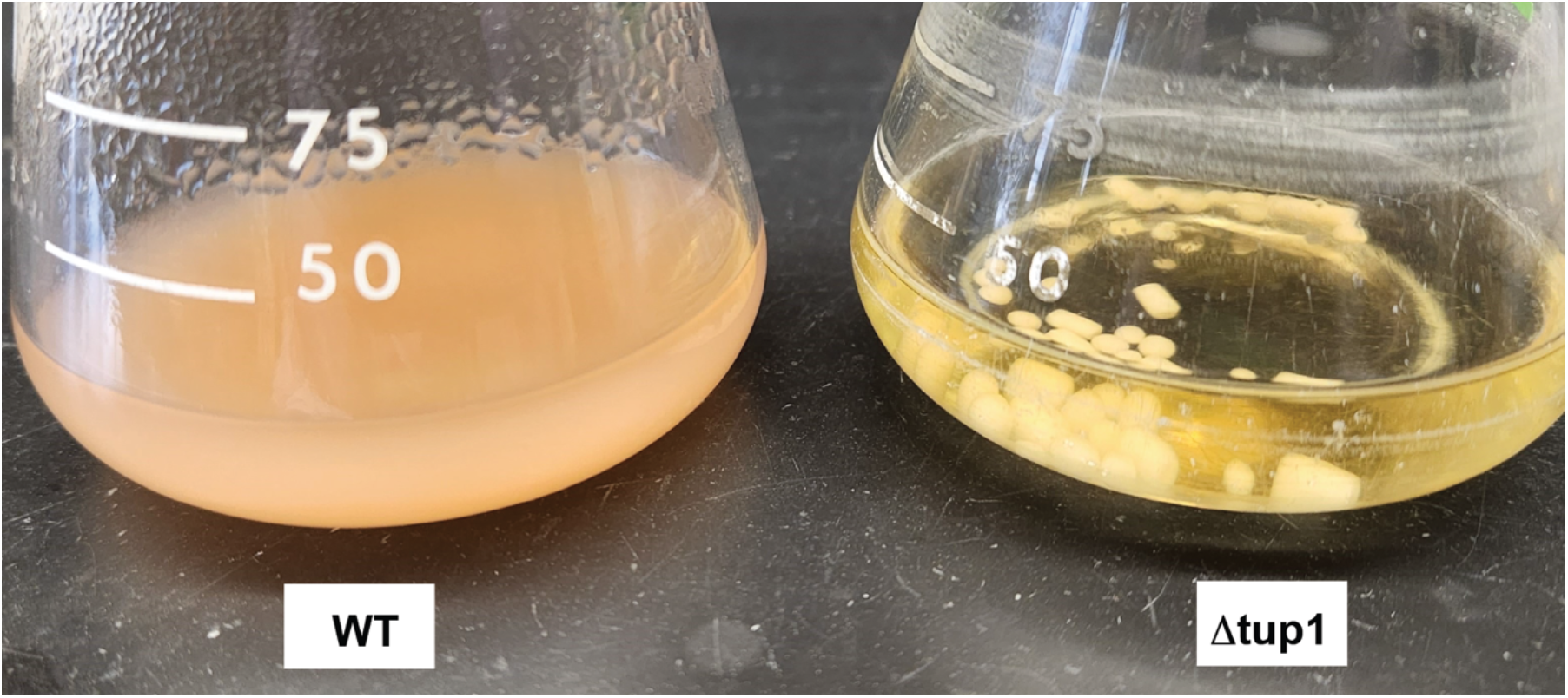
Deletion of Tup1 causes clumping of yeast cells. WT (left) and Δtup1 (right) yeast grown to log phase in YPD.

**Supplementary Figure 2.**
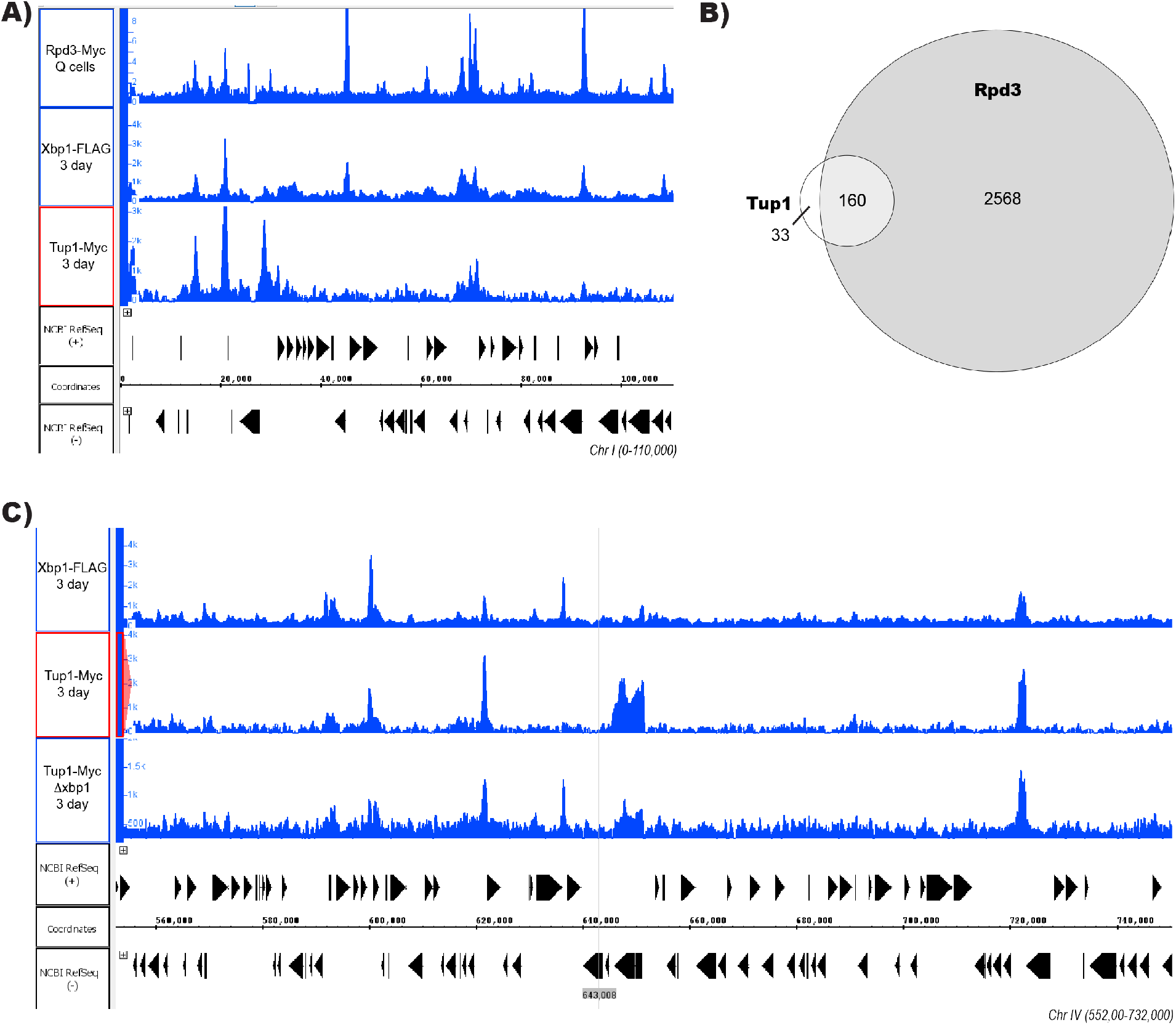
Rpd3, Xbp1, and Tup1 Colocalize in Glucose-Starved Yeast. (A) Integrated Genome Browser (IGB) tracks representative of ChIP-Seq for Rpd3-Myc in quiescent cells (Q cells), Xbp1-FLAG in stationary phase (3 day), and Tup1-Myc in stationary phase (3 day). Rpd3-Myc data is from McKnight, Boerma et al. 2015. (**B**) Venn diagram comparing peaks from Rpd3-Myc ChIP of purified Q cells and Tup1-Myc ChIP of 3-day cultures. **(C)** Integrated Genome Browser (IGB) tracks representative of ChIP-Seq in stationary phase (3 day) for Xbp1-FLAG, Tup1-Myc, and Tup1-Myc in a Δxbp1 background.

## Notes

https://www.ncbi.nlm.nih.gov/projects/geo/query/acc.cgi?acc=GSE210934

### Competing Interest Statement

The authors have declared no competing interest.

### Summary of Updates

The title has been changed for clarity, and minor changes have been made to the text, mostly grammatical.

