## Supplemental Table 1 for "Tup1 is Critical for Transcriptional Repression in Quiescence in *S. cerevisiae*"

| ALT_ID | CONSENSUS | TP | TP% | FP | FP% | ENR_RATIO | SCORE_THR | PVALUE | LOG_PVALUE | EVALUE | LOG_EVALUE | QVALUE | LOG_QVALUE |
|---|---|---|---|---|---|---|---|---|---|---|---|---|---|
| Urc2p | HCGGAGWTA | 57 | 29.23 | 12 | 6.15 | 4.46 | 13 | 3.330E-08 | -17.22 | 0.0000243 | -10.62 | 0.0000134 | -11.22 |
| Rap1p | ACACCCAYACAYYY | 64 | 32.82 | 16 | 8.21 | 3.82 | 12 | 5.430E-08 | -16.73 | 0.0000398 | -10.13 | 0.0000134 | -11.22 |
| Tod6p | ADCGCGATGAGSTNHNV | 28 | 14.36 | 1 | 0.51 | 14.5 | 13 | 7.890E-08 | -16.36 | 0.0000577 | -9.76 | 0.0000134 | -11.22 |
| Dot6p | ANGANGCGATGAGSTGH | 24 | 12.31 | 0 | 0 | 25 | 16 | 8.090E-08 | -16.33 | 0.0000592 | -9.73 | 0.0000134 | -11.22 |
| Mot3p | TAGGTA | 65 | 33.33 | 18 | 9.23 | 3.47 | 12 | 1.920E-07 | -15.47 | 0.000141 | -8.87 | 0.0000254 | -10.58 |
| Urc2p | CNTCGGAGATAWTT | 57 | 29.23 | 15 | 7.69 | 3.63 | 12 | 5.610E-07 | -14.39 | 0.000411 | -7.8 | 0.0000619 | -9.69 |
| Met32p | ANTGTGGCGB | 53 | 27.18 | 14 | 7.18 | 3.6 | 11 | 1.460E-06 | -13.44 | 0.00107 | -6.84 | 0.000138 | -8.89 |
| Rox1p | TCTATTGTTTCCC | 54 | 27.69 | 15 | 7.69 | 3.44 | 8.8 | 2.160E-06 | -13.05 | 0.00158 | -6.45 | 0.000179 | -8.63 |
| Ste12p | TTGAAACAATGAAACG | 50 | 25.64 | 13 | 6.67 | 3.64 | 8.4 | 2.430E-06 | -12.93 | 0.00178 | -6.33 | 0.000179 | -8.63 |
| Stp4p | DNNNNTTCAGCCGYACBAGNG | 70 | 35.9 | 25 | 12.82 | 2.73 | 11 | 3.830E-06 | -12.47 | 0.0028 | -5.88 | 0.000253 | -8.28 |
| Tec1p | ACATTCTTACATTCTT | 64 | 32.82 | 22 | 11.28 | 2.83 | 7.5 | 5.630E-06 | -12.09 | 0.00412 | -5.49 | 0.000339 | -7.99 |
| Azf1p | AAAAAGAAA | 60 | 30.77 | 20 | 10.26 | 2.9 | 12 | 7.180E-06 | -11.84 | 0.00526 | -5.25 | 0.000396 | -7.83 |
| Matalpha2p | ATTGTT | 76 | 38.97 | 30 | 15.38 | 2.48 | 8.9 | 8.230E-06 | -11.71 | 0.00603 | -5.11 | 0.000419 | -7.78 |
| YGR067C | GHGGGG | 75 | 38.46 | 30 | 15.38 | 2.45 | 9.6 | 1.170E-05 | -11.35 | 0.0086 | -4.76 | 0.000556 | -7.5 |
| Upc2p | CTCGTWTAG | 83 | 42.56 | 36 | 18.46 | 2.27 | 11 | 1.800E-05 | -10.92 | 0.0132 | -4.33 | 0.000795 | -7.14 |
| Srd1p | CNNTTTGTAGATCYCAH | 72 | 36.92 | 29 | 14.87 | 2.43 | 10 | 1.960E-05 | -10.84 | 0.0143 | -4.25 | 0.000809 | -7.12 |
| Mot3p | WAGGTA | 67 | 34.36 | 26 | 13.33 | 2.52 | 11 | 2.140E-05 | -10.75 | 0.0157 | -4.15 | 0.000835 | -7.09 |
| Pdr1p | TCCGCGGA | 48 | 24.62 | 15 | 7.69 | 3.06 | 11 | 2.880E-05 | -10.46 | 0.0211 | -3.86 | 0.000953 | -6.96 |
| Pdr3p | TCCGCGGA | 48 | 24.62 | 15 | 7.69 | 3.06 | 11 | 2.880E-05 | -10.46 | 0.0211 | -3.86 | 0.000953 | -6.96 |
| Rap1p | GGRTGTACGG | 48 | 24.62 | 15 | 7.69 | 3.06 | 10 | 2.880E-05 | -10.46 | 0.0211 | -3.86 | 0.000953 | -6.96 |
| Ste12p | ATGAAACAATGAGACA | 26 | 13.33 | 4 | 2.05 | 5.4 | 14 | 3.950E-05 | -10.14 | 0.0289 | -3.54 | 0.00115 | -6.77 |
| Msn1p | AATGTCCH | 54 | 27.69 | 19 | 9.74 | 2.75 | 12 | 3.980E-05 | -10.13 | 0.0291 | -3.54 | 0.00115 | -6.77 |
| YKL222C | BAACGGARATAANNC | 54 | 27.69 | 19 | 9.74 | 2.75 | 12 | 3.980E-05 | -10.13 | 0.0291 | -3.54 | 0.00115 | -6.77 |
| Tec1p | TTCTCACATTCTTC | 71 | 36.41 | 30 | 15.38 | 2.32 | 9.1 | 4.720E-05 | -9.96 | 0.0345 | -3.37 | 0.00129 | -6.65 |
| Zap1p | TTCTTTATGGT | 45 | 23.08 | 14 | 7.18 | 3.07 | 14 | 4.880E-05 | -9.93 | 0.0357 | -3.33 | 0.00129 | -6.65 |
| Hap1p | TAWCTCCG | 55 | 28.21 | 20 | 10.26 | 2.67 | 12 | 5.110E-05 | -9.88 | 0.0374 | -3.29 | 0.0013 | -6.64 |
| Gat3p | AAATBRGATCTACAAGCTG | 81 | 41.54 | 38 | 19.49 | 2.1 | 9.8 | 8.830E-05 | -9.34 | 0.0646 | -2.74 | 0.00216 | -6.14 |
| Tye7p | TGCRTCACGTGAYGCNNC | 22 | 11.28 | 3 | 1.54 | 5.75 | 13 | 1.000E-04 | -9.21 | 0.0732 | -2.61 | 0.00236 | -6.05 |
| YML081W | CCCCDCH | 107 | 54.87 | 57 | 29.23 | 1.86 | 10 | 1.110E-04 | -9.1 | 0.0816 | -2.51 | 0.00254 | -5.97 |
| Ste12p | CGTTTCA | 68 | 34.87 | 30 | 15.38 | 2.23 | 13 | 1.280E-04 | -8.96 | 0.0937 | -2.37 | 0.00282 | -5.87 |
| Yrm1p | ACGGAAAT | 66 | 33.85 | 29 | 14.87 | 2.23 | 12 | 1.510E-04 | -8.8 | 0.11 | -2.2 | 0.00322 | -5.74 |
| Yox1p | TWAATTR | 29 | 14.87 | 7 | 3.59 | 3.75 | 7.4 | 2.080E-04 | -8.48 | 0.152 | -1.88 | 0.00423 | -5.47 |
| Arg81p | AAGTGCAACTGACTGCGA | 25 | 12.82 | 5 | 2.56 | 4.33 | 7.6 | 2.110E-04 | -8.47 | 0.154 | -1.87 | 0.00423 | -5.47 |
| Sfl1p | THNHDNATAGAAGAAATAWDW | 32 | 16.41 | 9 | 4.62 | 3.3 | 13 | 2.910E-04 | -8.14 | 0.213 | -1.55 | 0.00567 | -5.17 |
| Stp4p | TGCGCTABC | 72 | 36.92 | 35 | 17.95 | 2.03 | 11 | 3.620E-04 | -7.93 | 0.265 | -1.33 | 0.00684 | -4.99 |
| Hcm1p | ATMAACAA | 63 | 32.31 | 29 | 14.87 | 2.13 | 10 | 3.970E-04 | -7.83 | 0.291 | -1.24 | 0.00731 | -4.92 |
| Stp3p | GCTAGCGCA | 55 | 28.21 | 24 | 12.31 | 2.24 | 12 | 4.800E-04 | -7.64 | 0.351 | -1.05 | 0.00859 | -4.76 |
| Lys14p | CNCAAAATTCCGBGCGNT | 47 | 24.1 | 19 | 9.74 | 2.4 | 12 | 5.470E-04 | -7.51 | 0.4 | -0.92 | 0.00953 | -4.65 |
| Abf1p | RTCRYYY | 51 | 26.15 | 22 | 11.28 | 2.26 | 10 | 6.690E-04 | -7.31 | 0.49 | -0.71 | 0.0114 | -4.48 |
| Fzf1p | YGSMNMCTATCAYTTYY | 40 | 20.51 | 15 | 7.69 | 2.56 | 11 | 7.060E-04 | -7.26 | 0.517 | -0.66 | 0.0117 | -4.45 |
| Pdr8p | TCCGHGGA | 88 | 45.13 | 49 | 25.13 | 1.78 | 9.2 | 9.120E-04 | -7 | 0.668 | -0.4 | 0.0147 | -4.22 |
| Rds2p | ANNNNGAAAAYCCGAGNNNNT | 55 | 28.21 | 26 | 13.33 | 2.07 | 11 | 1.240E-03 | -6.69 | 0.909 | -0.1 | 0.0196 | -3.93 |
| Gsm1p | ANCTCCG | 65 | 33.33 | 34 | 17.44 | 1.89 | 9.5 | 1.810E-03 | -6.32 | 1.32 | 0.28 | 0.0262 | -3.64 |
| Mig1p | ATTTTGCGGGG | 58 | 29.74 | 29 | 14.87 | 1.97 | 10 | 1.820E-03 | -6.31 | 1.33 | 0.29 | 0.0262 | -3.64 |

| | | | | | | | | | | | | | |
|---|---|---|---|---|---|---|---|---|---|---|---|---|---|
| Mig2p | ATTTTGCGGGG | 58 | 29.74 | 29 | 14.87 | 1.97 | 10 | 1.820E-03 | -6.31 | 1.33 | 0.29 | 0.0262 | -3.64 |
| Mig3p | ATTTTGCGGGG | 58 | 29.74 | 29 | 14.87 | 1.97 | 10 | 1.820E-03 | -6.31 | 1.33 | 0.29 | 0.0262 | -3.64 |
| Gcr1p | CWTCC | 52 | 26.67 | 25 | 12.82 | 2.04 | 9 | 2.000E-03 | -6.21 | 1.46 | 0.38 | 0.0282 | -3.57 |
| Cep3p | YTCGGAAA | 78 | 40 | 44 | 22.56 | 1.76 | 12 | 2.090E-03 | -6.17 | 1.53 | 0.43 | 0.0288 | -3.55 |
| Cup2p | HTHNNGCTGD | 27 | 13.85 | 9 | 4.62 | 2.8 | 12 | 2.490E-03 | -5.99 | 1.83 | 0.6 | 0.0337 | -3.39 |
| Stb5p | CGGTGTTA | 30 | 15.38 | 11 | 5.64 | 2.58 | 10 | 2.780E-03 | -5.89 | 2.03 | 0.71 | 0.0368 | -3.3 |
| Rlm1p | CTAWWWWTAG | 39 | 20 | 17 | 8.72 | 2.22 | 10 | 3.060E-03 | -5.79 | 2.24 | 0.81 | 0.0397 | -3.23 |
| Tbf1p | ARCCCTAA | 64 | 32.82 | 35 | 17.95 | 1.81 | 11 | 3.420E-03 | -5.68 | 2.5 | 0.92 | 0.0435 | -3.13 |
| YKL222C | AACGGARAT | 73 | 37.44 | 42 | 21.54 | 1.72 | 11 | 3.730E-03 | -5.59 | 2.73 | 1.01 | 0.0466 | -3.07 |
| Arg80p | CCTCTAAAGG | 51 | 26.15 | 26 | 13.33 | 1.93 | 9.2 | 4.100E-03 | -5.5 | 3 | 1.1 | 0.0503 | -2.99 |
| Srd1p | GTAGWTC | 42 | 21.54 | 20 | 10.26 | 2.05 | 9.6 | 4.800E-03 | -5.34 | 3.51 | 1.26 | 0.0578 | -2.85 |
| Mcm1p | TTACCNAATTNGGTAA | 110 | 56.41 | 72 | 36.92 | 1.52 | 4 | 4.960E-03 | -5.31 | 3.63 | 1.29 | 0.0578 | -2.85 |
| Ixr1p | KTTSAAYKGTTYASA | 105 | 53.85 | 68 | 34.87 | 1.54 | 6.6 | 4.970E-03 | -5.3 | 3.64 | 1.29 | 0.0578 | -2.85 |
| Abf1p | AGCCGTAAATAGTTATCTTCCAAG | 33 | 16.92 | 14 | 7.18 | 2.27 | 0.15 | 5.130E-03 | -5.27 | 3.76 | 1.32 | 0.0586 | -2.84 |
| Aft2p | CGCACCC | 101 | 51.79 | 66 | 33.85 | 1.52 | 9.4 | 6.670E-03 | -5.01 | 4.88 | 1.59 | 0.0749 | -2.59 |
| Yap1p | ASGTAATT | 108 | 55.38 | 72 | 36.92 | 1.49 | 10 | 7.240E-03 | -4.93 | 5.3 | 1.67 | 0.0799 | -2.53 |
| Nrg2p | HAGGGTCB | 50 | 25.64 | 27 | 13.85 | 1.82 | 11 | 7.970E-03 | -4.83 | 5.83 | 1.76 | 0.0865 | -2.45 |
| Cad1p | TTACGTAAT | 152 | 77.95 | 109 | 55.9 | 1.39 | 7.5 | 8.230E-03 | -4.8 | 6.02 | 1.8 | 0.0879 | -2.43 |
| Pdr3p | VCGGAA | 47 | 24.1 | 25 | 12.82 | 1.85 | 8.8 | 8.600E-03 | -4.76 | 6.3 | 1.84 | 0.0894 | -2.41 |
| Dal81p | GAAAATTGCGTT | 55 | 28.21 | 31 | 15.9 | 1.75 | 10 | 8.780E-03 | -4.74 | 6.43 | 1.86 | 0.0894 | -2.41 |
| Dal82p | GAAAATTGCGTT | 55 | 28.21 | 31 | 15.9 | 1.75 | 10 | 8.780E-03 | -4.74 | 6.43 | 1.86 | 0.0894 | -2.41 |
| Met32p | TGTGGCDB | 132 | 67.69 | 93 | 47.69 | 1.41 | 8.9 | 9.450E-03 | -4.66 | 6.92 | 1.93 | 0.0948 | -2.36 |
| Rds1p | CCBBTCGGCCGAAVDNCD | 89 | 45.64 | 58 | 29.74 | 1.53 | 7.2 | 1.000E-02 | -4.61 | 7.32 | 1.99 | 0.0988 | -2.31 |
| Rlm1p | TAWWWWTAGM | 81 | 41.54 | 52 | 26.67 | 1.55 | 9.2 | 1.110E-02 | -4.51 | 8.09 | 2.09 | 0.103 | -2.27 |
| Met32p | ABTGTGGC | 67 | 34.36 | 41 | 21.03 | 1.62 | 10 | 1.120E-02 | -4.49 | 8.22 | 2.11 | 0.103 | -2.27 |
| Mig1p | TTATTTCTGGGG | 108 | 55.38 | 74 | 37.95 | 1.45 | 6.4 | 1.130E-02 | -4.48 | 8.28 | 2.11 | 0.103 | -2.27 |
| Mig2p | TTATTTCTGGGG | 108 | 55.38 | 74 | 37.95 | 1.45 | 6.4 | 1.130E-02 | -4.48 | 8.28 | 2.11 | 0.103 | -2.27 |
| Mig3p | TTATTTCTGGGG | 108 | 55.38 | 74 | 37.95 | 1.45 | 6.4 | 1.130E-02 | -4.48 | 8.28 | 2.11 | 0.103 | -2.27 |
| Mot3p | AAGGWT | 50 | 25.64 | 28 | 14.36 | 1.76 | 11 | 1.140E-02 | -4.48 | 8.32 | 2.12 | 0.103 | -2.27 |
| Msn1p | AATGTCC | 39 | 20 | 20 | 10.26 | 1.9 | 12 | 1.190E-02 | -4.43 | 8.69 | 2.16 | 0.106 | -2.24 |
| Stb5p | CGGKGTTATA | 73 | 37.44 | 46 | 23.59 | 1.57 | 9.9 | 1.210E-02 | -4.41 | 8.89 | 2.19 | 0.107 | -2.23 |
| Rdr1p | TNNGNCNTGCGGAWATNNNNC | 60 | 30.77 | 36 | 18.46 | 1.65 | 12 | 1.280E-02 | -4.36 | 9.39 | 2.24 | 0.111 | -2.2 |
| Rph1p | AANNNDAWTTAGGGGKGNA | 84 | 43.08 | 55 | 28.21 | 1.52 | 9.5 | 1.290E-02 | -4.35 | 9.41 | 2.24 | 0.111 | -2.2 |

**Supplementary Table 1: Motif Analysis of Tup1 ChIP in Stationary Phase (3 day)**

TP = The number of **primary** sequences matching the motif / the number of primary sequences (the percentage of primary sequences matching the motif)

TP% = The percentage of primary sequences matching the motif with scores greater than or equal to the optimal match score threshold

FP = The number of **control** sequences matching the motif / the number of primary sequences (the percentage of control sequences matching the motif)

FP% = The percentage of control sequences matching the motif with scores greater than or equal to the optimal match score threshold

ENR_RATIO = The relative enrichment ration of the motif in the primary vs. control sequences, defined as

Ratio = ((TP+1)/(NPOS+1)) / ((FP+1)/(NNEG+1)),

where NPOS is the number of primary sequences in the input, and NNEG is the number of control sequences in the input

SCORE_THR = The match score threshold giving the optimal *p*-value. This is the score threshold used by SEA to determine the values of “TP” and “FP”
